# A yeast FRET biosensor enlightens cAMP signalling

**DOI:** 10.1101/831354

**Authors:** Dennis Botman, Tom G. O’Toole, Joachim Goedhart, Frank J. Bruggeman, Johan H. van Heerden, Bas Teusink

## Abstract

The cAMP-PKA signalling cascade in budding yeast regulates adaptation to changing environments. We developed yEPAC, a FRET-based biosensor for cAMP measurements in yeast. We used this sensor with flow cytometry for high-throughput single cell-level quantification during dynamic changes in response to sudden nutrient transitions. We found that the characteristic cAMP peak differentiates between different carbon source transitions, and is rather homogenous among single-cells, especially for transitions to glucose. The peaks are mediated by a combination of extracellular sensing and intracellular metabolism. Moreover, the cAMP peak follows Weber’s law; its height scales with the relative, and not the absolute, change in glucose. Lastly, our results suggest that the cAMP peak height conveys information about prospective growth rates. In conclusion, our yEPAC-sensor makes possible new avenues for understanding yeast physiology, signalling and metabolic adaptation.

## Introduction

*Saccharomyces cerevisiae*, or budding yeast, is a unicellular organism that lives in continuously changing environments to which it has to adequately adapt to stay competitive. To do so, yeast cells sense changes in nutrient availability and generate signals that they use as cues to adapt their physiological behaviour such as cellular metabolism and growth. For yeast, the most preferred carbon source is glucose, and it has evolved various signalling pathways responsive to its concentration^1–3^. One of these pathways is the cAMP-PKA pathway. Activation of cAMP-PKA occurs when derepressed cells are transitioned to an environment containing an abundant fermentable carbon source. This results in a transient increase of cAMP on a short timescale (i.e. seconds-minutes) and subsequent relaxation to an elevated steady level^4,5^. Activation of the cAMP-PKA pathway occurs via two distinct routes. First, import and metabolism of fermentable sugars activates Ras, stimulated by intracellular acidification^4,6–15^. Ras, in turn, activates the adenylate cyclase Cyr1 to produce cAMP. Second, the G-protein coupled receptor Gpr1 senses extracellular glucose and activates Cyr1 via Gpa2 (a G_α_ protein)^4,6,10,16–23^. Increased cAMP levels lead to activation of PKA, by causing dissociation of the regulatory subunit Bcy1 from the PKA subunits^24–26^. Activated PKA inhibits the stress-related transcription factors Msn2, Msn4 and the Rim15 protein kinase^27–30^. Moreover, PKA induces trehalose and glycogen breakdown^31–33^ and increases levels of the glycolytic activator fructose-2,6-bisphosphate^34–36^. Altogether, activation of the cAMP-PKA pathway induces a shift from a slow growth or stress-resistant physiological state to a fast growing fermentative one.

A rise in cAMP is transient, due to its rapid degradation by phosphodiesterase 1 and 2^37–39^. Moreover, the signalling cascade itself is inhibited via feedback inhibition through active PKA (which inhibits various cAMP signalling components). Additionally, cAMP signalling is inhibited by Ira1, Ira2 (which inhibits Ras) and by Rgs2 (which inhibits Gpa2) and these inhibitions also give rise the transient nature of the response^37,38,40–44^. A few studies suggest that the glycolytic intermediate fructose-1,6-bisphosphate is an activator of Ras and determines the basal cAMP levels^6,10,45^.

Although much progress has been made on cAMP-PKA signalling in yeast, various questions still remain. For the most part, characterisations were performed using solely glucose (or fructose) as fermentable carbon source. Therefore, cAMP responses to many other carbon sources or to stress-conditions are still largely unexplored. This is mainly because cAMP determination through conventional assay kits is rather labour intensive. Since only a few conditions are generally studied, input-output characterisations of cAMP-PKA signalling are scarce. It is also still unknown whether heterogeneity occurs in single-cell cAMP responses; Cell-to-cell heterogeneity is especially relevant for industrial bioprocessing where glucose-signalling heterogeneity can affect industrial efficiency^46–49^.

To address these questions, we developed a genetically-encoded biosensor for cAMP. We adapted an EPAC-based FRET sensor originally developed for mammalian cells^50,51^ for use in budding yeast. The resulting yeast-EPAC (yEPAC) sensor contains a FRET pair which is optimal for yeast, measures cAMP with high selectivity and shows a high FRET ratio change. This enables convenient intracellular measurements of cAMP in living single-yeast cells for the first time. We characterised cAMP responses at the single-cell level and in response to various nutrient transitions. Furthermore, we used flow-cytometry to quantify cellular heterogeneity in cAMP-dynamics during carbon source transitions. Combined, the obtained cAMP measurements with the new biosensor revealed several novel insights, including a strong dependence of cAMP peak height on the added carbon source and pre-growth conditions. Moreover, the use of yEPAC showed us that, against a fermentative background, the amplitude of the cAMP response (peak height) is a measure for the relative change (i.e. fold change) in glucose concentration, but against a respiratory background, peak height upon sugar addition appears to predict the extent of fermentative growth.

## Material and methods

### Fluorescent protein plasmids construction

The FRET-pairs mCherry-T2A-mTurquoise2 (mTq2), tagRFP-T2A-mTq2, tagRFPT-T2A-mTq2 and tdTomato-T2A-mTq2 in pDRF1-GW were previously constructed^52^. mCherry-mTq2, tagRFP-mTq2 and tagRFPT-mTq2 in a clontech-style C1 mammalian expression vector were obtained from Mastop et al.^53^, digested using NheI and NotI (New England Biolabs, Ipswich, Massachusetts, USA), and ligated with T4 ligase (New England Biolabs) in the yeast expression vector pDRF1-GW ^52^, digested with the same restriction enzymes.

mTq2 in pDRF1-GW was created by performing a PCR, using KOD polymerase (Merck-Millipore, Burlington, Massachusetts, USA) on mTq2-C1 using forward primer 5’-AGGTCTATATAAGCAGAGC-3’ and reverse primer 5’-TAGCGGCCGCTTACTTGTACAGCTCGTCCATG-3’. Next, the product and pDRF1-GW were digested with NheI and NotI and the PCR product was ligated into pDRF1-GW using T4 ligase, generating mTq2 in pDRF1-GW.

### yEPAC construction

mTq2Δ-Epac (CD,ΔDEP)-cp173Venus-cp173Venus (Epac-SH188) was a kind gift of dr. Kees Jalink. A PCR with KOD polymerase was performed on tdTomato-C1, using forward primer 5’-TAGAGCTCATGGTGAGCAAGGGCGAGG-3’ and reverse primer 5’-GCGGCCGCTTACTTGTACAGCTCGTCCATGCCG-3’. Next, both the PCR product and Epac-SH188 were digested, using SacI and NotI (New England Biolabs). The PCR product was ligated into Epac-SH188 using T4 ligase, replacing cp173Venus-cp173Venus for tdTomato. The adapted sensor and pDRF1-GW were digested using NheI and NotI and the sensor was ligated into pDRF1-GW, generating yEPAC (mTurquoise2Δ-Epac(CD,ΔDEP)-tdTomato in pDRF1-GW).

### Yeast transformation

Strains used in this study are described in table 1. These strains were transformed exactly as described by Gietz and Schiestl^54^.

### In vitro characterisation

W303-1A WT cells transformed with pDRF1-GW and yEPAC were grown overnight at 200 rpm and 30°C in 1x yeast nitrogen base (YNB) medium without amino acids (Sigma Aldrich, Stl. Louis, MO, USA), containing 100 mM glucose (Boom BV, Meppel, Netherlands), 20 mg/L adenine hemisulfate (Sigma-Aldrich), 20 mg/L L-tryptophan (Sigma-Aldrich), 20 mg/L L-histidine (Sigma Aldrich) and 60 mg/L L-leucine (SERVA Electrophoresis GmbH, Heidelberg, Germany). The next day, cells were diluted and grown to an OD600 of approximately 3 in 50 mL of the same medium. Next, cells were kept on ice and washed twice in ice-cold 20 mL 0.01 M KH2PO4/K2HPO4 buffer at pH7 containing 0.75 g/L EDTA (AppliChem GmbH, Darmstadt, Germany). After the last wash step, cells were resuspended in 2 mL of 0.01 M KH2PO4/K2HPO4 buffer containing 0.75 g/L EDTA. Cells were washed twice in 1 mL of ice-cold 0.1 M KH2PO4/K2HPO4 buffer at pH7.4 containing 0.4 g/L MgCl2 (Sigma-Aldrich). Cells were transferred to screw cap tubes pre-filled with 0.75 grams of glass beads (425-600 μm) and lysed using a FastPrep-24 5G (MP Biomedicals, Santa Ana, CA, USA) with 8 bursts at 6 m/s and 10 seconds per burst. Afterwards, the lysates were centrifuged for 15 minutes at 21000 g and the cell-free extracts were snap-frozen in liquid nitrogen and stored at −80°C for later use.

Per sample, 5 wells of a black 96-well microtitre plate (Greiner Bio-One) were filled with 40 μL of cell-free extract. Fluorescence spectra were recorded after successive additions of cAMP (Sigma-Aldrich) using a CLARIOstar platereader (BMG labtech, Ortenberg, Germany). Spectra were recorded with 430/20 nm excitation and 460-660 nm emission (10 nm bandwidth). Fluorescence spectra were corrected for background fluorescence (by correcting for fluorescence of W303-1A WT expressing the empty pDRF1-GW plasmid) and FRET ratios were calculated by dividing donor over acceptor fluorescence. The data was fitted to the Hill equation (equation 1)^51^, with cAMP denoting the cAMP concentration, K_d_ the dissociation constant, and n the Hill-coeffecient.

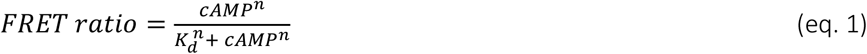

### ConA solution

Concanavalin A (ConA) was prepared as described by Hansen et al., 2015^55^. In brief, 5 mg of ConA (Type IV, Sigma Aldrich) was dissolved in 5mL PBS at pH6.5, 40 mL H_2_O, 2.5mL of 1 M MnCl_2_ and 2.5 mL of 1 M CaCl_2_ and stored at −80°C.

### Microscopy

Strains used in this study are described in table 1. These strains (expressing yEPAC) were grown overnight at 200 rpm and 30°C in 1x YNB medium without amino acids, containing 20 mg/L adenine hemisulfate, 20 mg/L L-tryptophan, 20 mg/L L-histidine, 60 mg/L L-leucine and either 1% Ethanol (v/v, VWR International, Radnor, PA, United States of America), 1% glycerol (v/v, Sigma Aldrich), 100 mM pyruvate (Sigma Aldrich) or 111 mM galactose (Sigma Aldrich). Next, cells were diluted in the same medium and grown to an OD600 of maximally 1.5 and with minimal 5 cell divisions. The cultures were transferred to a 6-well microtitre plate containing cover slips pre-treated ConA to immobilize the cells. Afterwards, the coverslip was put in an Attofluor cell chamber (Thermofisher Scientific, Waltham, MA, USA) and 1 mL of fresh medium was added. Samples were imaged with a Nikon Ti-eclipse widefield fluorescence microscope (Nikon, Minato, Tokio, Japan) at 30°C equipped with a TuCam system (Andor, Belfast, Northern Ireland) containing 2 Andor Zyla 5.5 sCMOS Cameras (Andor) and a SOLA 6-LCR-SB power source (Lumencor, Beaverton, OR, USA). Fluorescent signals were obtained using a 438/24 nm excitation filter. The emission was separated by a 552 nm long-pass (LP) dichroic filter in a TuCam system. A 483/32 nm and 593/40 nm emission filter-pair was used for the detection of donor and acceptor emission, respectively (all filters from Semrock, Lake Forest, IL, USA). Perturbations were performed by adding 1x YNB medium containing the same amino acids as described before with 10x concentrated carbon source or KCl to the cell chamber to the desired concentration. Per condition, at least 2 biological replicates were obtained. Cells were segmented and fluorescence was measured with an in-house FiJi macro (NIH, Bethesda, MD, USA).

**Table 1.**
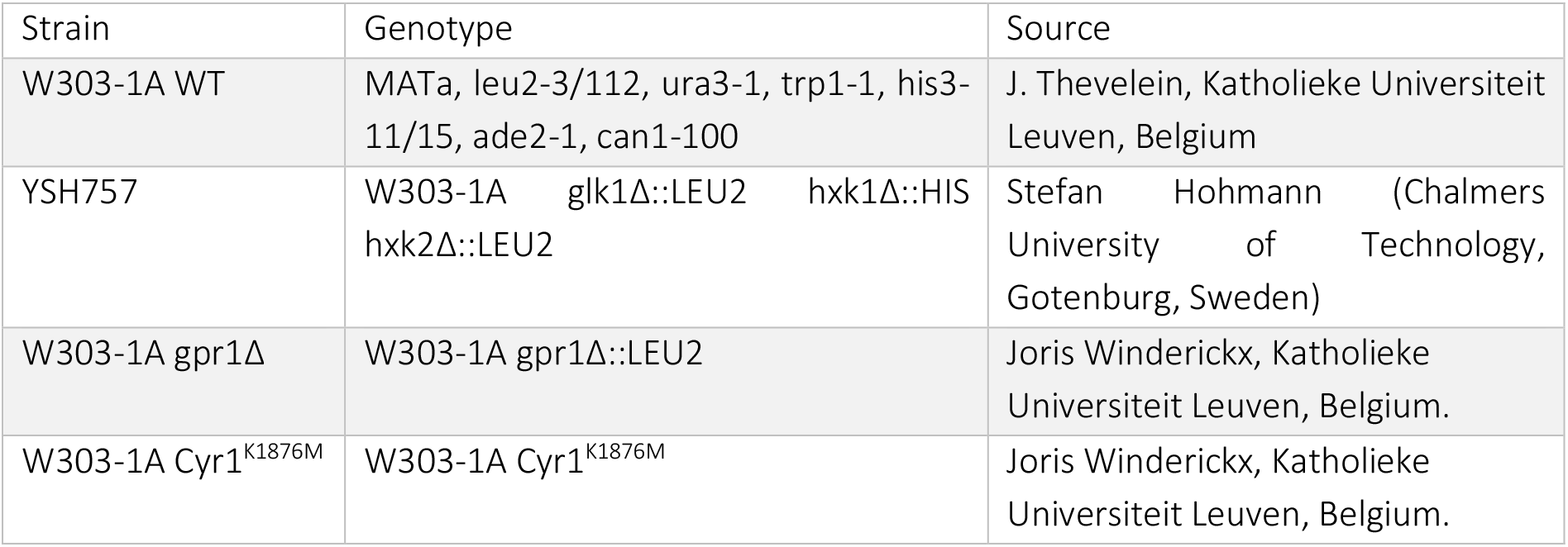
*S. cerevisiae* strains used in this paper.

### Growth experiments

Cells expressing pDRF1-GW and yEPAC were grown to midlog as described for microscopy with medium containing 1% ethanol. Next, cells were washed and resuspended to an OD600 of 1 with the same medium with the carbon source omitted. Cells were transferred to an OD of 0.05 in a 48-well microtitre plate containing 480 μL of fresh medium with either 0.1% ethanol, 10 mM galactose or 10 mM glucose. The cells were grown in a Clariostar plate reader at 30°C and 700 rpm orbital shaking. OD600 was measured every 5 minutes.

### Fluorescence lifetime imaging and spectral imaging

W303-1A WT cells expressing mCherry-mTq2, mCherry-T2A-mTq2, tagRFP-mTq2, tagRFP-T2A-mTq2, tagRFPT-mTq2, tagRFPT-T2A-mTq2, tdTomato-mTq2 and tdTomato-T2A-mTq2 were grown for at least 2 weeks on 2% agarose plates containing 6.8 g/L YNB without amino acids, 100mM glucose, 20 mg/L adenine hemisulfate, 20 mg/L L-tryptophan, 20 mg/L L-histidine and 60 mg/L L-leucine. Frequency domain FLIM was performed as described before^53^. Briefly, 18 phase images were obtained with a RF-modulated image intensifier (Lambert Instruments II18MD, Groningen, The Netherlands) set at a frequency of 75.1MHz coupled to a CCD camera (Photometrics HQ, Tucson, AZ, USA) as detector. mTq2 was excited using a directly modulated 442nm laser diode (PicoQuant, Berlin, Germany). Emission was detected using a 480/40 nm filter. The lifetimes were calculated based on the phase shift of the emitted light (τϕ). Per sample, 3 replicates were recorded.

Emission spectra of a donor-acceptor fusion protein or unfused equimolar expressed donor and acceptor were acquired as described previously^53^. In brief, excitation was at 436/20 nm and the emission was passed through a 80/20 (transmission/reflection) dichroic mirror and a 460 nm LP filter. Individual spectra were corrected for expression level by quantifying the intensity of the acceptor by excitation at 546/10 nm and detection with a 590 nm LP filter. Per sample, 3 replicates were measured.

### Flow cytometry

W303-1A strains expressing yEPAC, pDRF1-GW and mTq2 were grown as described for microscopy. Flow cytometry was performed using an BD INFLUX cell sorter (Becton Dickinson, Franklin Lakes, NJ, USA), with a 140 μm nozzle and a sheath pressure of 6 psi to run the samples. The sorter was equipped with a 200 mW Solid State 488 nm laser focused on pinhole 1, a 75 mW Solid State 561 nm laser focused on pinhole 3 with a laser delay of 18.17 μsec and a 100 mW Solid State 445 nm laser focused on pinhole 5 with a laser delay of 37.11 μsec. PMTs (photo multiplier tubes) for the 445 nm laser and the 488 nm laser were assimilated in trigon detector arrays that use serial light reflections – moving from the longest wavelengths to the shortest – to collect the dimmest emission signals first. The 445 nm trigon array was configured with a 610/20 nm bandpass filter in detector A and a 520/35 nm bandpass filter (preceded by a 502 nm LP filter) in detector B. The 488 trigon array was configured with a 610/20 nm bandpass (preceded by a 600LP) in detector A, a 530/40 nm bandpass (preceded by a 520 LP) in detector B and a 488/10 bandpass in detector C (SSC). PMTs for the 561 laser were assimilated in an octagon detector array. Acceptor emission was measured in detector D which was filtered with a 610/20 bandpass (preceded by a 600LP). Per condition, at least 2 biological replicates were obtained. All events were corrected for background fluorescence (median fluorescence of cells expressing pDRF1-GW), bleedtrough corrected (median fluorescence of cells expressing mTq2 only in the acceptor channel) and filtered for saturating or low fluorescence and scatter values. The effect of sensor expression on FRET ratios were calculated by plotting FRET ratios against tdTomato expression, obtained with the 561 nm laser and a 610/20 nm bandpass filter.

### pH sensitivity

Cells expressing yEPAC-R279L and mVenus-mTq2 were grown to an OD_600_ of maximally 1.5 in YNB medium containing 100 mM glucose. Cells were washed 3 times and resuspended in Citric Acid/Na_2_HPO_4_ buffer at various pH containing 2 mM of the ionophore 2,4-dinitrophenol to equilibrate pH levels. Afterwards, FRET ratios were recorded using a widefield microscope as described before.

### Data analysis

All data were analysed and visualized using R version 3.5.1 (R Foundation for Statistical Computing, Vienna, Austria). For analysis, moving and dead cells were manually removed. Additionally, cells with low fluorescence (i.e. below 50 A.U. fluorescence counts) were excluded. Next, acceptor fluorescence was corrected for bleedtrough (12% of total donor fluorescence) and FRET ratios were normalized to the mean FRET ratio before the perturbation (baseline). Dose-response kinetics were fitted using equation 2 with Peak_max_ denoted as the maximal peak height that can be obtained, glucose the amount of glucose pulsed and K_0.5_ the glucose amount that induces half the maximal peak height.

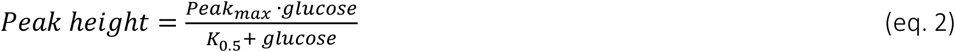

## Results

### Engineering of a versatile EPAC sensor for cAMP quantifications in yeast

In yeast, it is still a challenge to measure cAMP levels continuously in living single cells since a commendable cAMP FRET sensor for yeast is lacking and the current standard of cAMP determination still relies on population-averaged cAMP determination using cell extracts at single timepoints. Therefore, we took a mammalian-optimized EPAC sensor^56^ and replaced the tandem-cp173Venus acceptor with tdTomato since this fluorescent protein turned out to be a better acceptor in yeast with acceptable maturation, a higher photostability, brightness and pH-robustness compared to Venus (Fig. S1)^52,57^. Furthermore, tdTomato is a good acceptor for mTq2, with a FRET efficiency of 23%, a substantial sensitized emission and no effect of expression levels on FRET ratios (Fig. S1, table S1), making it very suitable for ratiometric fluorescence readouts. We named this sensor yEPAC.

*In vitro* calibration of yEPAC showed loss-of-FRET upon cAMP addition, and therefore, FRET ratios are presented as CFP/RFP ratios in this paper. We determined that yEPAC has a K_D_ of 4 μM for cAMP, which is slightly lower compared to the original sensor (Fig. 1A) and in the range of physiological cAMP levels in yeast^37,58–60^. We performed various control experiment to characterize the performance and potential of yEPAC. We confirmed that glucose addition to cells grown on a non-fermentable carbon source indeed gave a transient cAMP peak up to baseline normalized FRET values of 1.7 (Fig. 1B). Furthermore, the Cyr1^K1876M^ mutation and introduction of the R279L in de cAMP-binding domain showed a largely diminished cAMP response (Fig. 1B), as expected^61,62^. The small response of the R279L sensor variant is probably caused by osmotic changes, since addition of the non-metabolizable sugar sorbitol gave an identical response of this sensor (Fig. S1D). Importantly, the yEPAC sensor did not affect growth at various carbon sources (Fig. S2). Conversely, however, we did find a small growth rate effect on the FRET levels of the sensor (Fig. S3). This makes the sensor less suitable to compare basal cAMP levels at various growth rates. The sensor had improved temporal resolution compared to the conventionally used cAMP assay kits since we could record cAMP responses up to 15 minutes with a 3 second time interval (Fig. 1C, movie S1).

**Figure 1.**
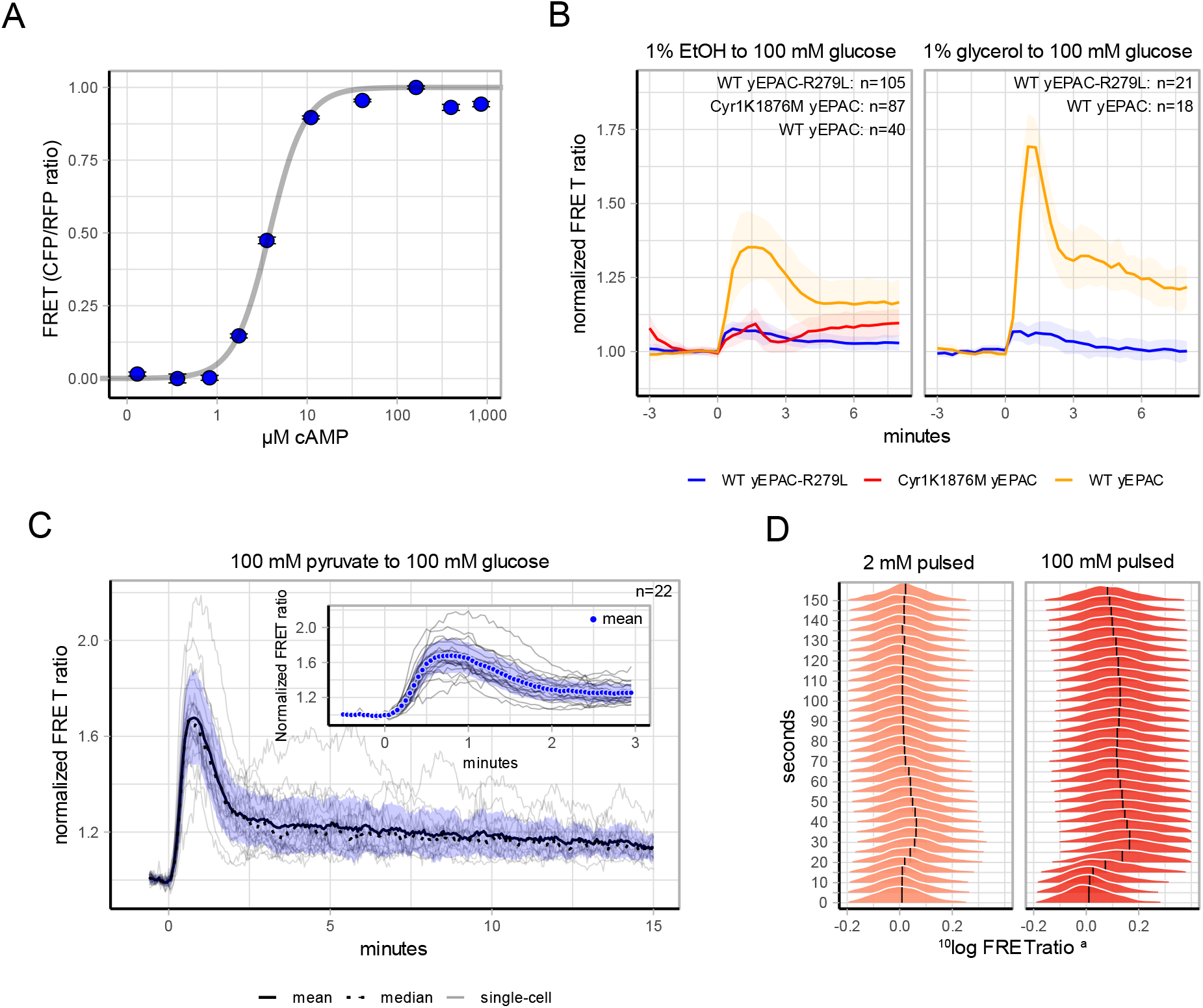
yEPAC characterisation. A) *In vitro* cAMP dose-response curve of yEPAC. Points indicate the mean FRET value of 5 replicates, error bars indicate SD. Solid line shows the Hill-fit (eq. 1). B) W303-1A WT cells expressing yEPAC or the non-responsive yEPAC-R279L and W303-1A cells that possessed the Cyr1^K1876M^ mutation were grown on 1% EtOH or 1% glycerol and pulsed with glucose at t=0 minutes. FRET signals were obtained and baseline normalized, lines show mean FRET ratios, shaded areas indicate SD. C) Pyruvate-grown W303-1A cells pulsed with 100 mM glucose at t=0 minutes. Inset shows the first 3 minutes after the pulse. FRET ratios are normalized to the baseline, solid lines show mean FRET ratios, dotted lines show median FRET ratios, grey lines show single-cell trajectories, shaded areas indicate SD. D) Dynamic frequency distribution of FRET values after 2 and 100 mM glucose addition, respectively. W303-1A WT cells were pre-grown on 1% EtOH, a baseline was recorded (not shown in graph) and a glucose pulse was added at 0 seconds. Timepoints were binned for every 5 seconds. Percentages are v/v, abbreviations: EtOH, ethanol, ^a^FRET ratios were baseline-normalized.

yEPAC can also be used in flow cytometry which provides a useful complement to microscopy, as it allows for hundreds to thousands of single-cells to be sampled per second. However, this technique cannot measure FRET in the same cells over time. We tested this method with additions of 2 or 100 mM glucose to ethanol grown cells (Fig. 1D). We obtained FRET ratios of 300-700 cells per second to determine cAMP responses. The dynamics were comparable to the microscopy-obtained data (Fig. 1D and S4). Of note, we did not observe non-responders, also not by the addition of 2 mM glucose.

In summary, our developed yEPAC sensor can be used to reliably measure cAMP in single yeast cells without adverse effects. Also, we show that our sensor can be used with flow cytometry, in addition to the conventional microscopy readouts, expanding its utility.

### cAMP peak heights follow the Weber-Fechner law

After using saturating glucose amounts (Figs. 1B, C & D), we studied how cAMP levels would change in response to lower amounts of glucose. We pulsed ethanol-grown W303-1A cells with glucose ranging from 0 to 50 mM. Normalized peak heights of these transitions showed a saturating dose-response with a K_0.5_ of 3.0 mM and a maximal peak height of 1.38 normalized FRET values (Fig. 2A).

**Figure 2.**
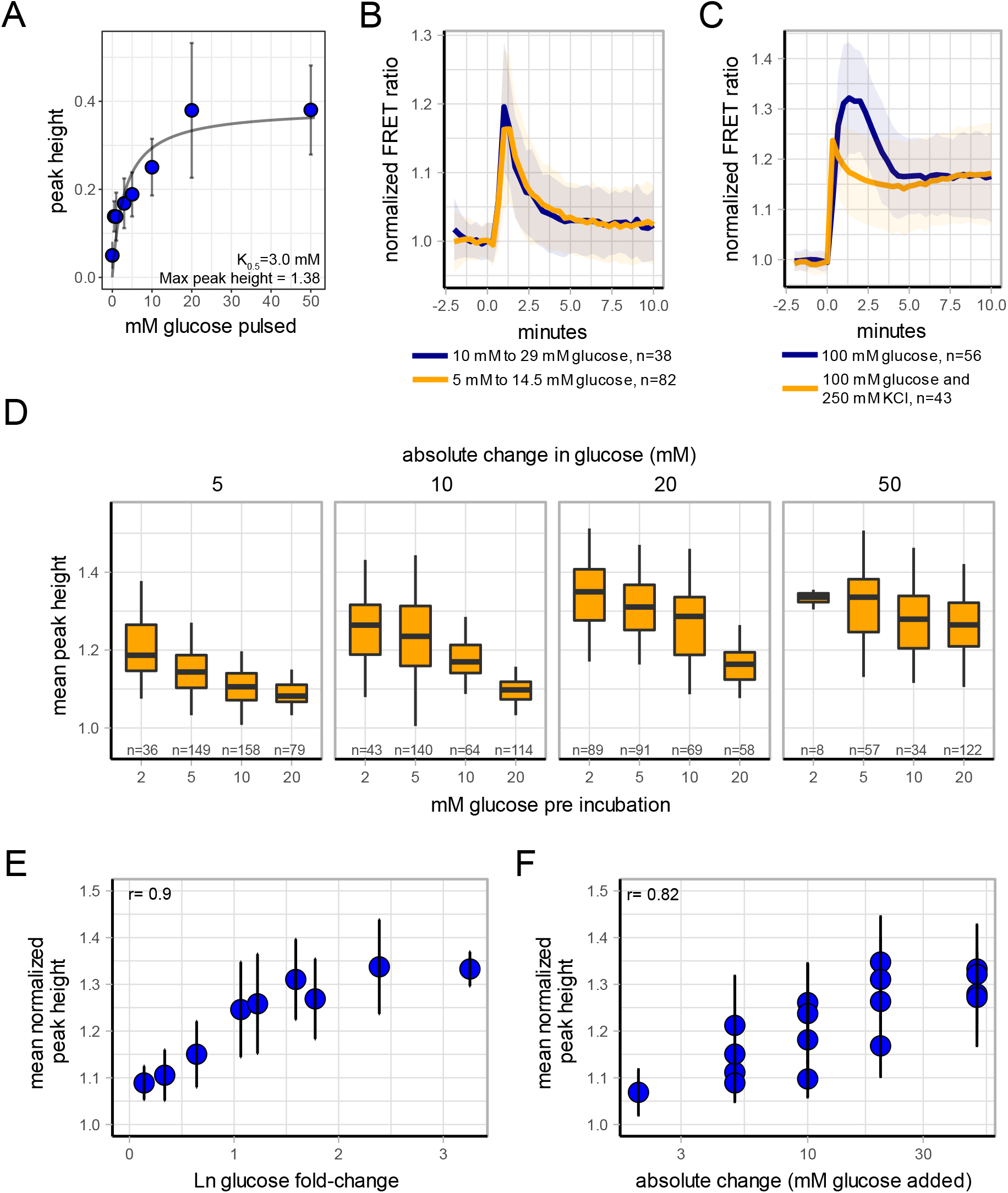
Dose-response and Weber-Fechner law experiments of cAMP. A) W303-1A cells expressing yEPAC were grown on 1% ethanol and various glucose concentrations were pulsed. Fitting peak heights versus the glucose concentration shows saturation kinetics with a K_0.5_ of 3.0 mM. Dots indicate mean value, error bars indicate SD, grey line shows the fit using eq. 2. B) W303-1A cells expressing yEPAC were pre-incubated at either 5 or 10 mM glucose and a 2.9-fold change of glucose was performed. Lines show mean response, shaded areas indicate SD. C) W303-1A cells expressing yEPAC were grown on 1% EtOH and either 100 mM glucose or 100 mM glucose with 250 mM KCl was added. Lines show mean responses, shaded areas indicate SD. D) Population response of cAMP of cells expressing yEPAC. Cells were incubated at various initial amounts of glucose (depicted below each graph) and various amounts of glucose were added (depicted above each graph). Dots show mean populations response, error bars indicate SD. E) Peak heights plotted against the log fold-change of various glucose transitions, dots indicate mean value, error bars indicate SD. F) Peak heights plotted against the absolute glucose added for transitions, dots indicate mean value, error bars indicate SD.

The dose-response data were generated against a background of zero glucose, and indicated that yeast cells are able to detect small amounts of glucose. However, we hypothesized that any advantage cells may reap from responding to a change in sugar availability, will depend largely on the amount of sugar already in the environment (i.e. the background level). This means that a cell should respond to small glucose changes when glucose levels are low, but should be less sensitive to the same (absolute) changes when glucose levels are already high. We therefore tested whether the magnitude of the glucose-induced cAMP peak heights scale with the relative change (i.e. fold-change) of the glucose concentration instead of the absolute change, *i.e.* whether it obeys the Weber-Fechner law^63^. We incubated cells in media with various background concentrations of glucose and subsequently added various amounts of glucose (Figs. 2B, D, E, F). Indeed, we found comparable responses between transitions with the same relative change but with different absolute amounts of glucose pulsed (Fig. 2B). Because systems that detect relative changes add up inputs with positive and negative responses^64^, we also tested the application of two such inputs simultaneously. First, we identified salt stress as a negative input as this reduced cAMP levels transiently (Fig. S5). Indeed, addition of salt stress to cells reduces glucose-induced peak heights (Fig. 2C), which indicates that the cAMP peak height can measure relative glucose changes. Normalized peak heights decreased with increasing amounts of pre-incubated glucose, which shows that cells indeed change cAMP peak heights based on their background level (i.e. the pre-incubated glucose level). Moreover, the normalized peak heights relate better with the fold-change in glucose than with the absolute glucose change (Figs. 2E and 2F).

Next, we tested whether a fixed glucose fold-change pulse applied successively to the same population, elicits a similar peak response (Fig. 3). Indeed, we found that also in this case, the baseline-normalized peak height scales with the relative glucose change and not the absolute amount. (Fig. 3A, B). For individual cells we found only a weak correlation between normalized-peak heights of the first versus the second perturbation.(Fig. 3C). This indicates that at the single cell level, the relative change is rather noisy, where the population response is robust. However, an increased time between the fold-changes may improve Weber-Fechner law detection at single-cell level. In line with this, we found that perturbations within a shorter timescale (i.e. every 10 minutes) showed deteriorated Weber-Fechner law responses (Fig. S6). In these transitions, cells largely lose their ability to detect relative glucose changes. This suggests that the minimal time scales at which cells can adapt their glucose threshold is in the order of 20-30 minutes, and therefore the mechanism likely involves changes in protein expression.

**Figure 3.**
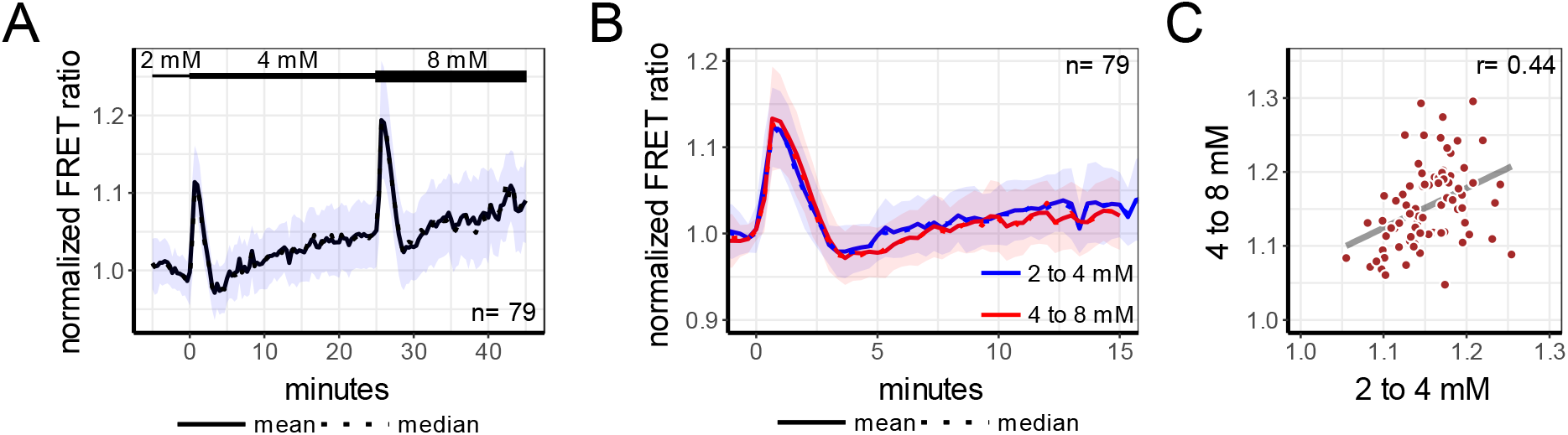
Weber-Fechner law of the same cells in time. A) W303-1A cells expressing yEPAC were pre-incubated on 2 mM glucose. Afterwards, cells were transitioned to 4 mM glucose at t=0 minutes and transitioned again to 8 mM at t=25 minutes, resulting in a two-fold change each time. Solid line shows the population mean response, dotted line shows median response, normalized to the first 5 minutes. B) Responses of the transitions performed in graph A, normalized to the last 3 frames before each transition. Solid line shows the population mean response, dotted line shows median response. Colour indicates the transition. C) Relation of normalized peak heights of single-cells between the first and second transitions. Shaded areas indicate SD. Dots represents single-cells, r-value shows the Spearman correlation coefficient (p < 0.01).

In conclusion, we show that cAMP responses are sensitive to glucose changes when cells reside in low glucose environments. In high glucose environment cAMP responses are rescaled, making the cAMP response relative to the current glucose levels cells are in.

### cAMP responses are carbon-source transition dependent

The cAMP signalling cascade is well known and characterised for its transitions from non-fermentable carbon sources to glucose. However, less data is available for other transitions. Therefore, we quantified the cAMP response for transitions between a variety of carbon sources. W303-1A WT cells were grown in medium containing 1% ethanol (v/v), 1% glycerol (v/v), 100 mM pyruvate or 111 mM galactose and subsequently pulsed with saturating amounts of either glucose, fructose, galactose, mannose or various oligosaccharides. This resulted in 18 different transitions for which the cAMP levels were monitored (Fig. 4).

**Figure 4.**
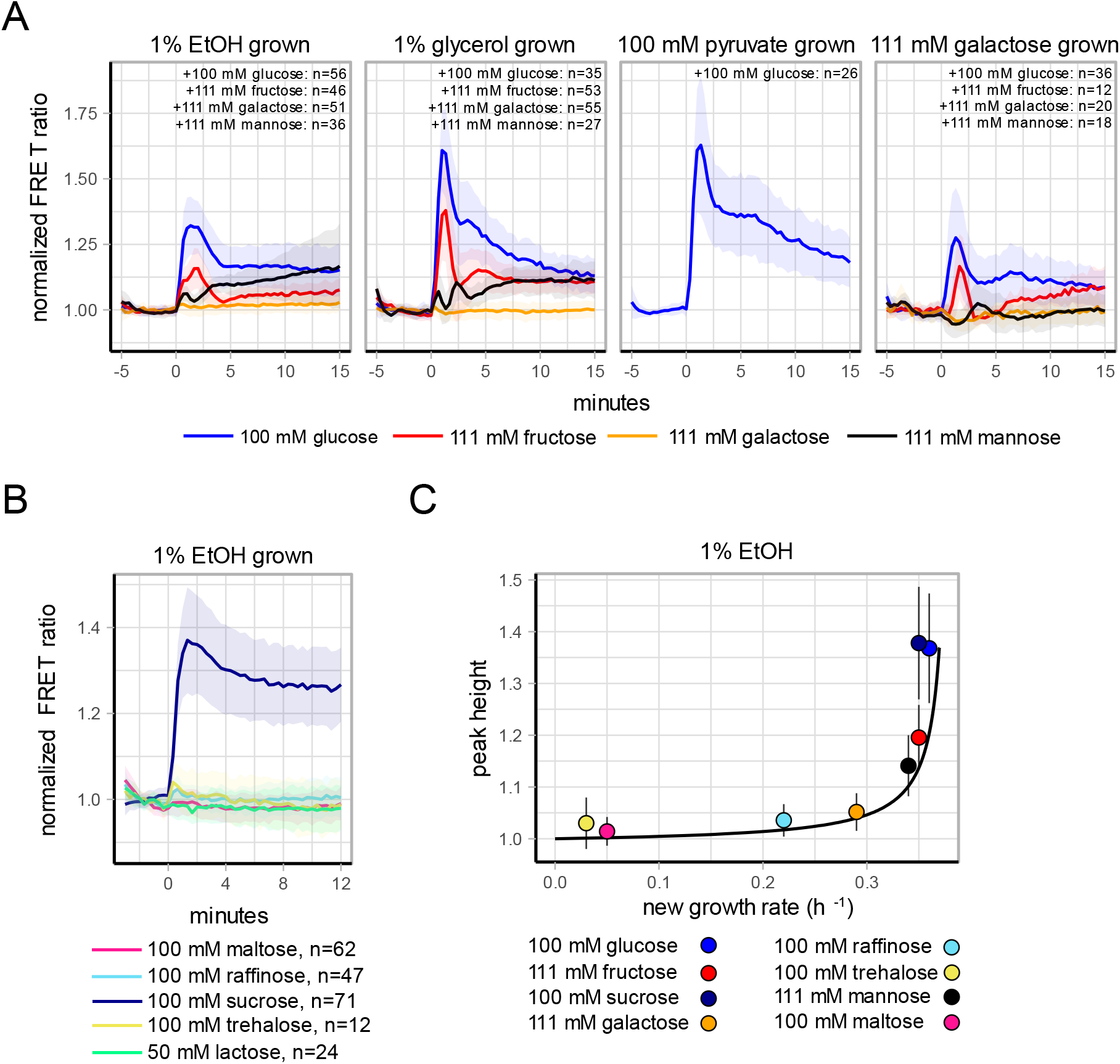
yEPAC FRET responses of various carbon transitions. A) Cells growing on various non-fermentative carbon sources (depicted above each graph) were pulsed with various sugars, depicted by line colour, at 0 minutes. Lines show mean FRET ratios, normalized to the baseline, shaded areas indicate SD. B) Cells growing on 1% ethanol were pulsed with various di- or trisaccharides and the FRET responses were recorded. Lines show mean FRET ratios, normalized to the baseline, shaded areas indicate SD, sugar was added at t=0 minutes. C) Peak heights of the transitions shown in A and B plotted against the maximal growth rate that can be obtained with the added carbon sources. Points show population mean, errorbars indicate SD, dotted line shows a glucose-dependent curve (obtained by plotting the dose response kinetics against Monod kinetics).

Among the added sugars, only sucrose, glucose and fructose induced clear cAMP peaks. Addition of sucrose gave the highest peak and fructose the lowest. Cells pulsed with galactose or mannose did not show a cAMP peak. Mannose is known as an antagonist of Gpr1^22,23^ and galactose is not a Gpr1 activator, which suggests that Gpr1 regulates the height of the cAMP peak. Noteworthy is the observation that mannose-pulsed cells did show increased cAMP levels after 15 minutes, even though an initial peak-response was absent. cAMP dynamics also depended on the pre-growth condition, with glycerol and pyruvate grown cells producing significantly higher cAMP peaks compared to EtOH and galactose grown cells (Wilcoxon test, p<0.01). In line with the dynamic flow-FRET results, we did not observe subpopulations or non-responders in most transitions (Figs. S7 and S8). Occasionally, we found for sugars other than glucose that some cells exhibited deviated cAMP dynamics compared to the population response. These cells show no cAMP peak or steadily increasing cAMP levels throughout the time-lapse recording. Therefore, cAMP signalling appears very robust for glucose transitions, but shows less robustness for other sugars.

Transitions from one primary carbon source to another can alter the maximal obtainable growth rate of cells. We wondered if cAMP peak heights could contain information about this new potential growth rate. Figure 4C indeed suggests such a relation: all data appear to lie on a curve. This curve fits very well with a glucose-dependent curve obtained when the dose response kinetics of figure 2A (max peak height 1.38 and K_0.5_ = 3.0 mM) is plotted against the growth rate inferred from published Monod kinetics with a maximal growth rate of 0.37 h^−1^ and a Ks of 0.1 mM^65^. Note that the cAMP peak height shows a sharp increase when a growth rate higher than 0.3 h^−1^ can be obtained, which is around the onset of overflow metabolism^66^.

In summary, our results show that the cAMP dynamics are context dependent, and that -at least for transitions from EtOH to sugars tested here- the peak height corresponds to the growth rate that can be achieved in the new environment.

### cAMP dynamics are affected by both Gpr1 signalling and sugar metabolism

Lastly, we examined which components of the signalling cascade affect the cAMP peak during transitions. cAMP signalling mutants were grown on medium containing 1% EtOH and pulsed with 2% glucose (Fig. 5). Deletion of either Gpr1, all three glucose phosphorylating enzymes (hxk1Δ, hxk2Δ, glk1Δ triple mutant) or the mutation in Cyr1 (Cyr1^K1876M^) affected the transient peak in cAMP. Noteworthy, the hxk1Δ, hxk2Δ, glk1Δ mutant still showed a clear cAMP peak (although decreased compared to WT). This mutant does not display transient intracellular acidification upon glucose addition (since glucose cannot be metabolized), indicating that cAMP peak generation does not solely rely on acidification, or metabolism for that matter. Deletion of another input via the membrane bound Gpr1-sensor, had a similar effect on the cAMP peak response. Since these two branches are known to regulate cAMP responses, we hypothesise that the residual cAMP response for both mutants are caused by the other cAMP signalling branch that is still functioning.

**Figure 5.**
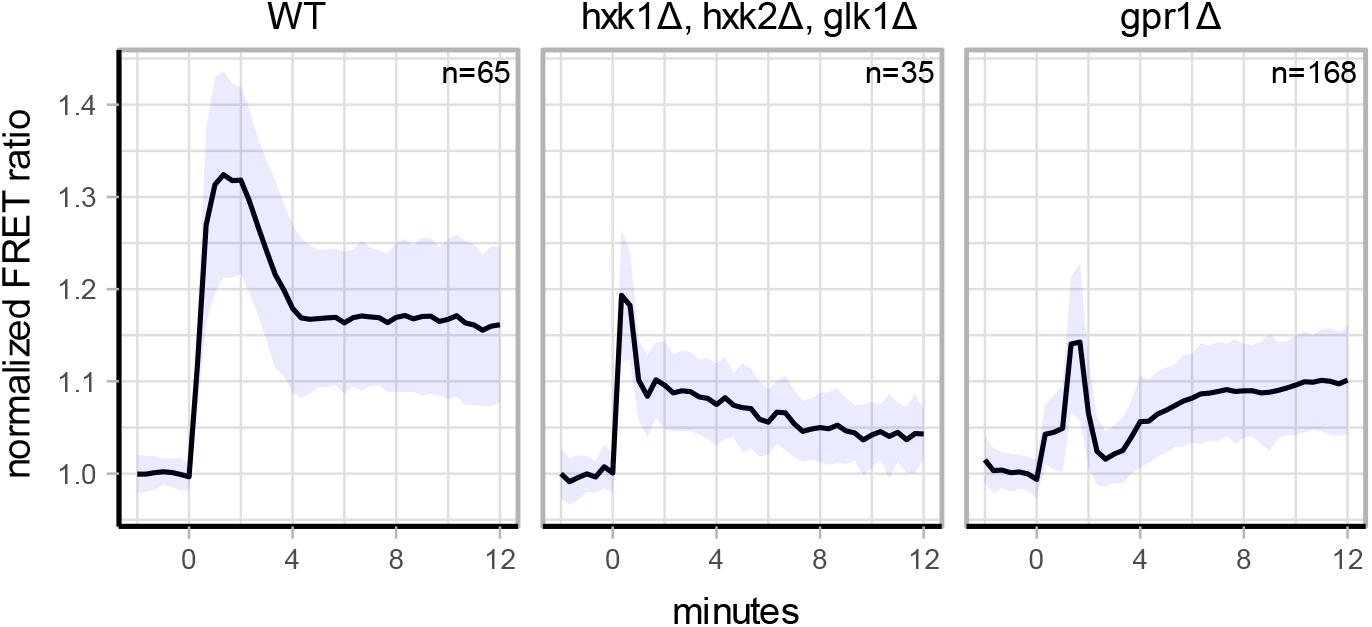
yEPAC FRET responses of various signalling mutants. The various cAMP signalling mutants in the W303-1A background were pulsed with 100 mM glucose at 0 minutes. Lines show mean FRET ratios, normalized to the baseline, shaded areas indicate SD.

## Discussion

We present a FRET-based biosensor for dynamic, single-cell cAMP detection in yeast. Although a different FRET-based cAMP EPAC biosensor for budding yeast was published recently^67,68^, we believe that our yEPAC has significant improvements. It shows a high FRET range, up to a normalized ratio change of 1.7 (Fig. 1B). Furthermore, the growth assays show that the sensor has no adverse effects on yeast physiology (Fig. S2). The slight difference in cAMP affinity of yEPAC compared to the original sensor can be caused by the different acceptor used, possibly changing conformation of the sensor slightly, or differences in characterisation such as the host species and the used buffers. Signalling mutants and the non-responsive yEPAC variant showed good cAMP selectivity of the sensor. However, we found a slight bias of basal FRET levels of yEPAC-R279L at various growth rates (Fig. S3). The origin of this bias is currently unknown and subject for future research and sensor improvements. The obtained flow-cytometry data gave a high temporal resolution compared to the conventional used cAMP assays. These results showed a clear secondary peak (Fig. S4) at high glucose concentrations. This peak was also present (e.g. Fig 4A), but not consistently observed, in the microscopy dataset. This oscillatory behaviour is in line with predictions from modelling efforts, but was not further explored in this study^69^.

Of note, flow cytometry, nor microscopy showed clear non-responders for glucose transitions (Fig. S7 and S8). However, we observed some heterogeneity in transitions with sugars that do not activate Gpr1^23^. Therefore, we hypothesise that this heterogeneity occurs from variation in the metabolism of the sugar, as observed before with carbon-source transitions^57,70^. These results show that the cAMP signalling cascade is robust, in contrast to what was found for pH^70^ and recently for intracellular ATP dynamics as well^57^. Apparently, as shown earlier^71^, signalling glucose or metabolising it are different challenges to yeast cells.

In nature, yeast cells likely encounter large fluctuations in glucose availability, ranging from complete absence to saturating amounts of glucose. Until now, it was unknown how cAMP signalling reacts to a glucose increase when glucose is already present in the environment. We tested these transitions and found that cAMP peak heights seem to measure glucose changes relative to the background level of glucose, a property known as the Weber-Fechner law. Our analyses to test for Weber-Fechner law assume that each cell performs baseline-normalization. This could give an extra benefit by reducing variability among cells^64,72–74^. However, cells need time to establish a baseline between successive glucose-additions, since shortening the period between glucose additions no longer gave scaling of the cAMP peak height with the relative fold change (Fig. S6).

In order to reliably test for Weber-Fechner law, cAMP levels should be below saturation values of the yEPAC sensor. We validated this by comparing the normalized peak heights in response to saturating glucose amount (i.e. 50 mM glucose), when cells are pre-incubated at 2, 5, 10 and 20 mM glucose (Fig. 2D). Normalized peak heights were comparable, indicating that cAMP levels did not yet saturated the sensor. Furthermore, pre-incubation with 2 to 20 mM glucose did not aspecifically affect the yEPAC sensor as the yEPAC-R279L hardly shows any response after pulsing with 100 mM glucose (Fig. S9), which is at least 5 times more than used for Weber-Fechner law characterisation. Our data therefore indicates that the cAMP pathway in yeast cells can detect (and adapt to) small glucose additions when glucose levels were low, but does not signal this change when glucose levels are high already. When cells are already fully fermentative, growing on glucose, the peak cannot indicate an increase in growth rate, but rather may signal how much sugar is present and whether or not cells should keep investing in fermentation and ribosomal biosynthesis.

This is different when we added different carbon sources to fully respiratory, ethanol pre-grown cells. At this background, a consistent relation between peak height and the prospective growth rate on the pulsed sugar was found, for different sugars, and fitted the predicted growth rate for different glucose concentrations, based on Monod growth kinetics. This suggests that the peak height informs about growth. The cAMP signalling cascade is generally considered to mediate a switch to a fermentative (i.e. high growth rate) mode. This was consistent with our data, where cAMP peak height increased sharply around the onset of fermentation, *i.e.* under conditions that generate a growth rate higher than 0.3 h^−1^, Fig. 4C). However, our results also show that cells without a clear cAMP peak (e.g. mannose pulsed cells) still obtain a high growth rate, without displaying a transient cAMP peak. Also the industrially important CEN.PK strain that has the K1876M mutation in Cyr1^61^ does not show a peak but does ferment. So although we find clear and interesting relationships, their functional implications remain to be fully elucidated.

The cAMP responses to various other sugar transitions and in signalling mutants show that cAMP dynamics are complex and highly context dependent. We could infer several features about cAMP-signalling from these data.

First, nine sugars were tested and only sucrose (giving the highest peak) and its breakdown products, glucose and fructose induced a cAMP peak. It is remarkable that yeast developed a signalling cascade for only these sugars. On the other hand, sucrose is often the end-product of plant photosynthesis and therefore one of the most abundant sugars in plants^75^. In nature, yeast resides on plants or fruit and sensing extracellular sucrose to consume conceivably improves yeast’s fitness.

Second, the data indicate that a cAMP peak is generated when either the initial metabolism of a sugar is sufficiently rapid or Gpr1 is activated. Combined activation is needed to achieve a maximal peak response. We found that fructose does not interact with Gpr1, but does induce a cAMP peak. In the case of fructose the peak is lower than peaks induced by the Gpr1 agonists sucrose or glucose, pointing to the amplifying effect of combined activation.

In stark contrast to the peak responses triggered by sucrose, glucose and fructose, mannose, which is an antagonist of Gpr1, does not show any cAMP peak. One explanation for the absence of a peak is signal-dampening through Gpr1 inhibition. Another explanation is that mannose gets transported much more slowly, since the hexose transporters have a lower V_max_ and higher K_m_ for mannose compared to glucose and fructose, which likely reduces the initial uptake rate of mannose^76,77^. Still, we found that mannose does trigger a gradual increase in cAMP levels shortly after its addition, which could indicate Ras activation through an increased glycolytic flux^45^. Accordingly, addition of galactose, which does not interact with Gpr1 and is not considered a rapidly-fermentable carbon source, does not induce a cAMP peak and does not yet show signs of a gradual increase in cAMP levels shortly after its addition. Indeed, cells growing on ethanol or glycerol are not immediately ready to metabolize galactose^78^, and we expect the cAMP levels to gradually rise with the induction of galactose metabolism. A differential response to mannose by galactose- or ethanol-grown cells further underscores the effect of pre-growth conditions on the ability of cells to sense and respond to sudden sugar transitions. Mannose did not show the gradual increase in cAMP levels in galactose-grown cells as it did in ethanol grown cells. Galactose growth suppresses the expression of various high-affinity hexose transporters, such as HXT6 and 7, compared to growth on ethanol or glycerol^79–81^. Mannose uptake rate is therefore expected to be much lower in galactose grown cells than ethanol or glycerol grown cells, which may explain these observations. The response of glucose and fructose addition to galactose-grown cells are expected as galactose-grown have a higher capacity to metabolize these sugars, which induces ^7,57,77,82,83^. Finally, we confirm that the cAMP peaks clearly originate partly from both the metabolism of the sugar as well as the Gpr1 receptor, as described before^18,22,84^. In line with previous studies, our data indicates that intracellular acidification is not a requisite as the hxk1Δ, hxk2Δ, glk1Δ shows no intracellular acidification (due to the absence of sugar phosphorylation) and still shows cAMP production^6,15,84^.

Overall, yEPAC enabled us for the first time to investigate single-cell cAMP dynamics and elucidate conveniently various input-output relations during various carbon-source transitions. This gave important new insights: the normalized peak height seems to be a signal for future growth rate on the pulsed sugar and is only produced when cells should switch to fermentative growth. Possibly, the peak height functions as a switch for rewiring to fermentable metabolism.

## Acknowledgements

We thank Kees Jalink (Division of Cell Biology, The Netherlands Cancer Institute) for sharing mTurquoise2Δ-Epac(CD,ΔDEP)-cp173Venus-cp173Venus (Epac-SH188). We thank Juan Garcia Vallejo and Cora Chadick (Molecular Cell Biology & Immunology, VUmc) for his help with flow cytometry. We are grateful to Joris Winderickx (Functional Biology, KU Leuven) and Marco Siderius (Amsterdam Institute for Molecules, Medicines and Systems (AIMMS), Division of Medicinal Chemistry) for sharing W303-1A mutant strains. We thank Daan de Groot and Philipp Savakis for fruitful discussions.

## Material requests

Requests for the yEPAC sensor should be addressed to Kees Jalink (Division of Cell Biology, Netherlands Cancer Institute, Plesmanlaan 121, 1066CX Amsterdam, the Netherlands). Other material requests should be addressed to Bas Teusink

## Supplements

Movie S1. Ratiometric movie of W303-1A WT cells expressing yEPAC. Cells were grown on 100 mM pyruvate and 100 mM glucose was pulsed. Colour indicates FRET ratio, depicted by the calibration bar in the upper right.

**Figure S1.**
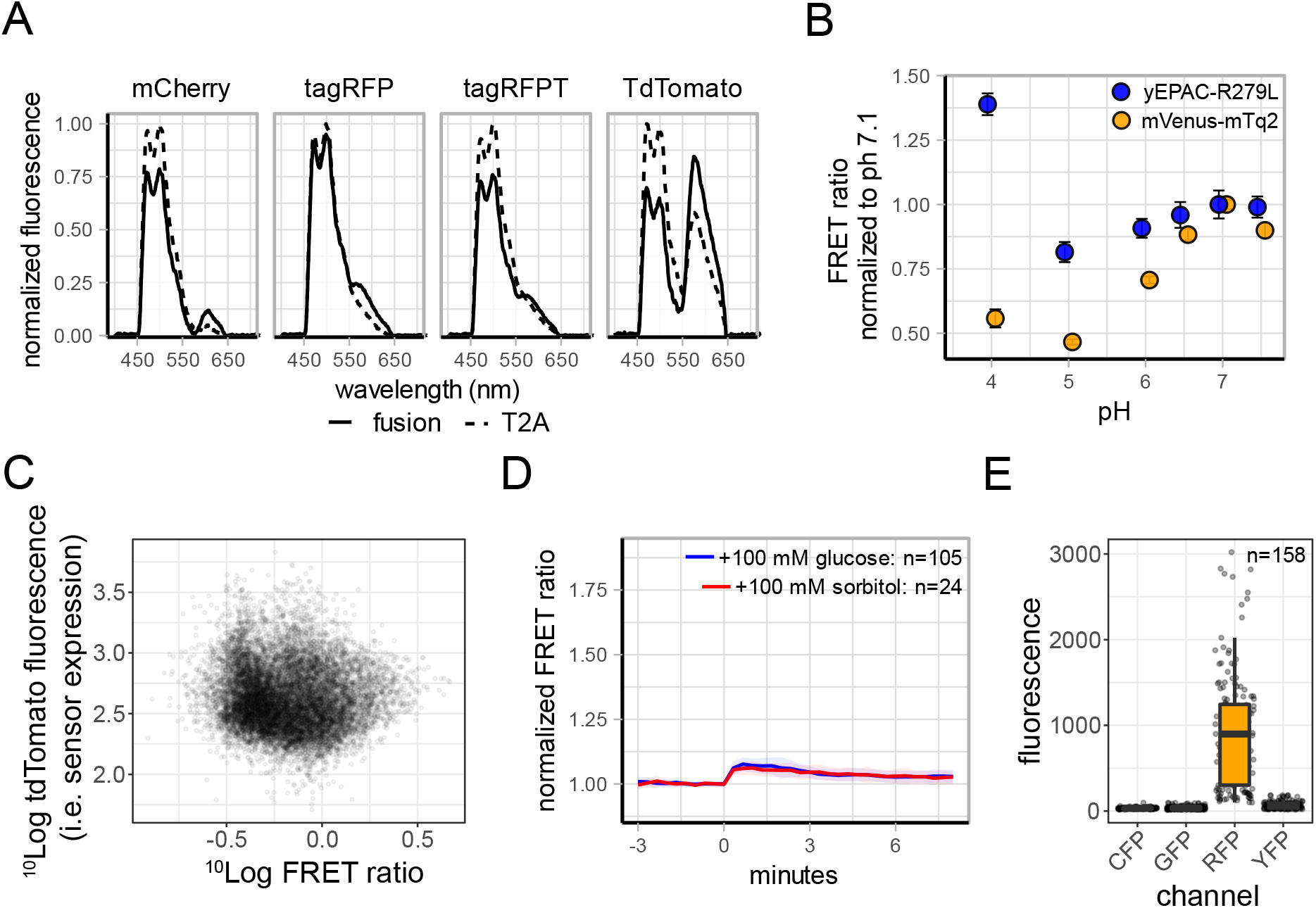
Characterisation of FRET pairs and yEPAC sensor. A) Spectra of various acceptors for mTq2. Solid lines show direct fusions of the depicted FP with mTq2. Dotted lines show spectra of the T2A fusion with mTq2, resulting in equimolar expression of cleaved mTq2 and acceptor, used as non-FRET control. Spectra of plated W303-1A colonies were recorded. B) pH sensitivity analyses of yEPAC-R279L and the mVenus-mTq2 FRET pair shows pH robustness up to pH 5. Cells expressing yEPAC-R279L and mVenus-mTq2 were grown to midlog in YNB medium containing 100 mM glucose. Cells were washed resuspended in Citric Acid/Na2HPO4 buffer containing 2 mM of the ionophore 2,4-dinitrophenol to equilibrate pH levels. Afterwards, FRET ratios were recorded using a widefield microscope. Points indicate mean FRET ratio, normalized to pH 7.1, errorbars indicate 95% CI. C) Expression of the yEPAC sensor (the fluorescence of tdTomato, measured by direct excitation) plotted against the FRET ratio show no clear relation between expression levels of the sensor and the measure FRET ratio. D) Response of yEPAC-R279L on the non-metabolizable sugar sorbitol. W303-1A WT cells expressing yEPAC-R279L were grown on 1% EtOH and pulsed with 100 mM sorbitol or 100 mM glucose at t=0 minutes. FRET signals were obtained and baseline normalized, lines show mean FRET ratios, shaded areas indicate SD. E) maturation characterisation of tdTomato. W303-1A WT cells expressing tdTomato were grown on 1% EtOH and CFP, GFP, RFP and YFP fluorescence was obtained. Points indicate single-cell fluorescence values. Boxplots indicate median values with quartiles, whiskers indicate largest and smallest observation at 1.5 times the interquartile range

**Table S1.** FRET efficiencies determined by frequency domain lifetime measurements of W303-1A colonies expressing the fusion constructs or mTq2. ^a^calculated lifetime of mTq2 determined by frequency domain. ^b^FRET efficiency calculated as (1-(Lifetime_donor+acceptor_/lifetimedonor))*100.

| Fusion | Lifetime<sup>a</sup> | FRET efficiency<sup>b</sup> |
| --- | --- | --- |
| mCherry-mTq2 | 3.29 ns | 12.5 |
| tagRFP-mTq2 | 3.73 ns | 0.8 |
| tagRFPT-mTq2 | 3.47 ns | 7.7 |
| TdTomato-mTq2 | 2.9 ns | 22.9 |
| mTq2 | 3.76 ns | n/a |

**Figure S2.**
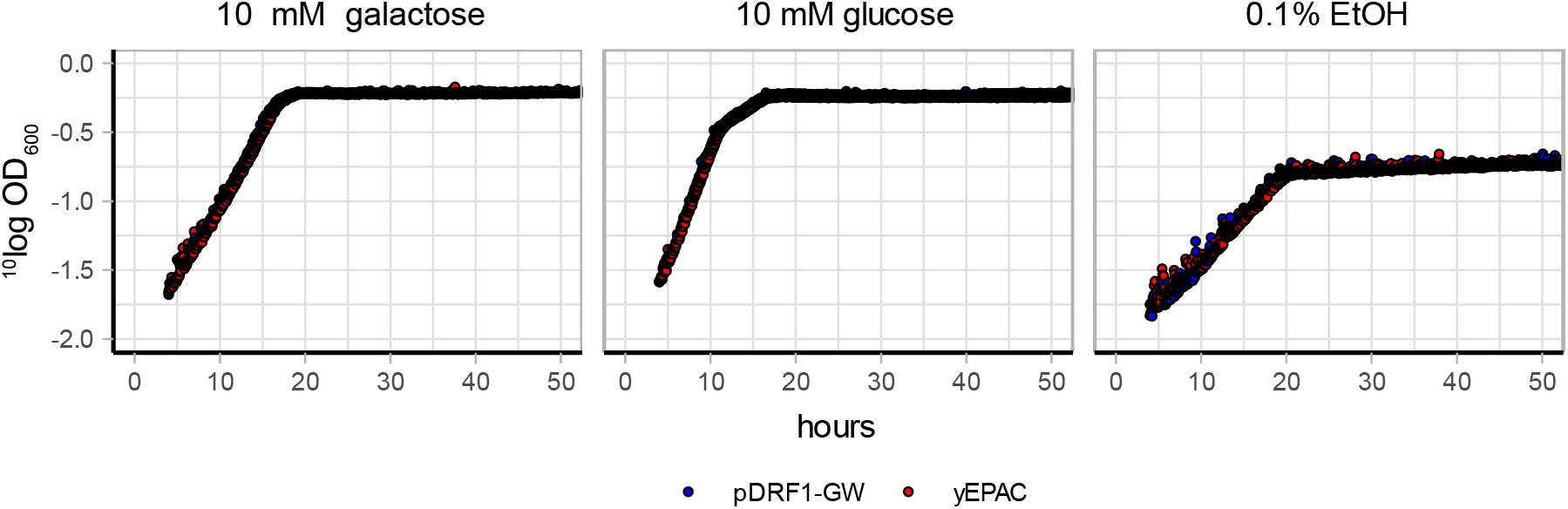
Growth curves of W303-1A WT cells expressing either yEPAC or the empty pDRF1-GW vector. Cells were grown to midlog in 1x YNB containing 1% EtOH, washed and resuspended in medium containing either 10 mM galactose, 10 mM glucose or 0.1% EtOH. Points indicate mean OD, color indicates the strain.

**Figure S3.**
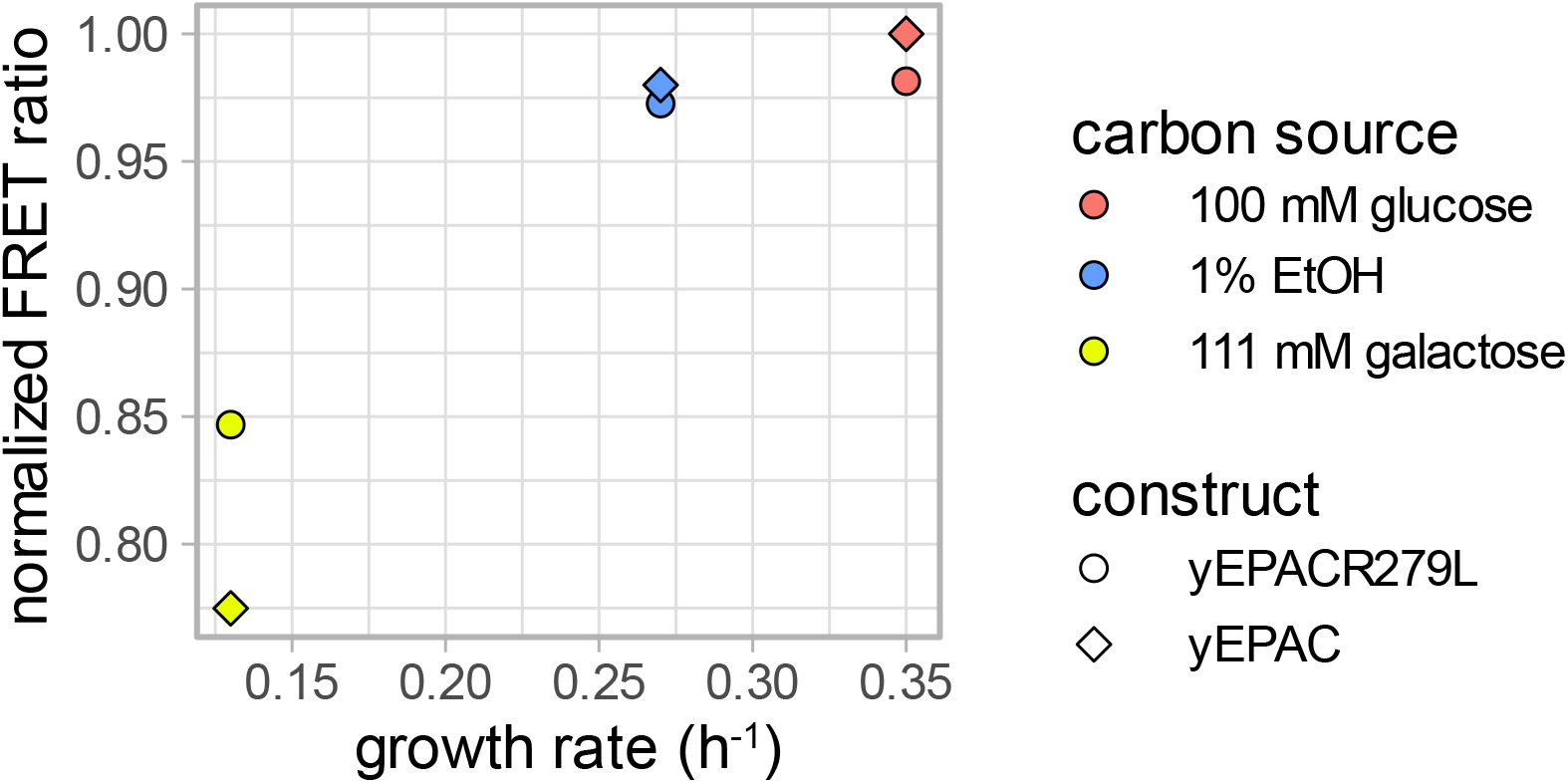
Baseline FRET ratios of W303-1A WT cells expressing yEPAC or the non-responding yEPAC-R279L shows a small baseline effect of growth rate on the FRET levels. Points indicate the median FRET level, colours indicate the carbon source cells grew on.

**Figure S4.**
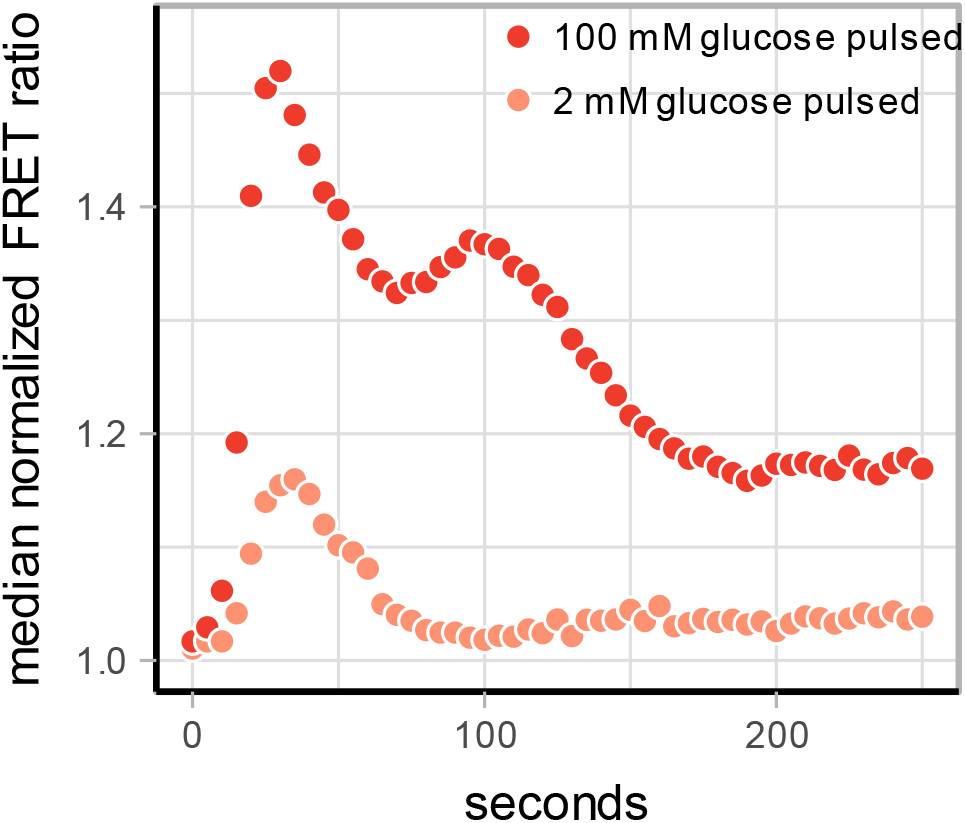
Dynamic flow-cytometry FRET measurements. B) W303-1A WT cells were grown on 1% EtOH, a baseline was recorded (not shown in graph) and a glucose pulse was added. The first obtained timepoint after glucose addition is set to t=0 seconds. Median FRET ratio for every binned timepoint are shown, depicted by the points. Point color indicate amount of glucose pulsed.

**Figure S5.**
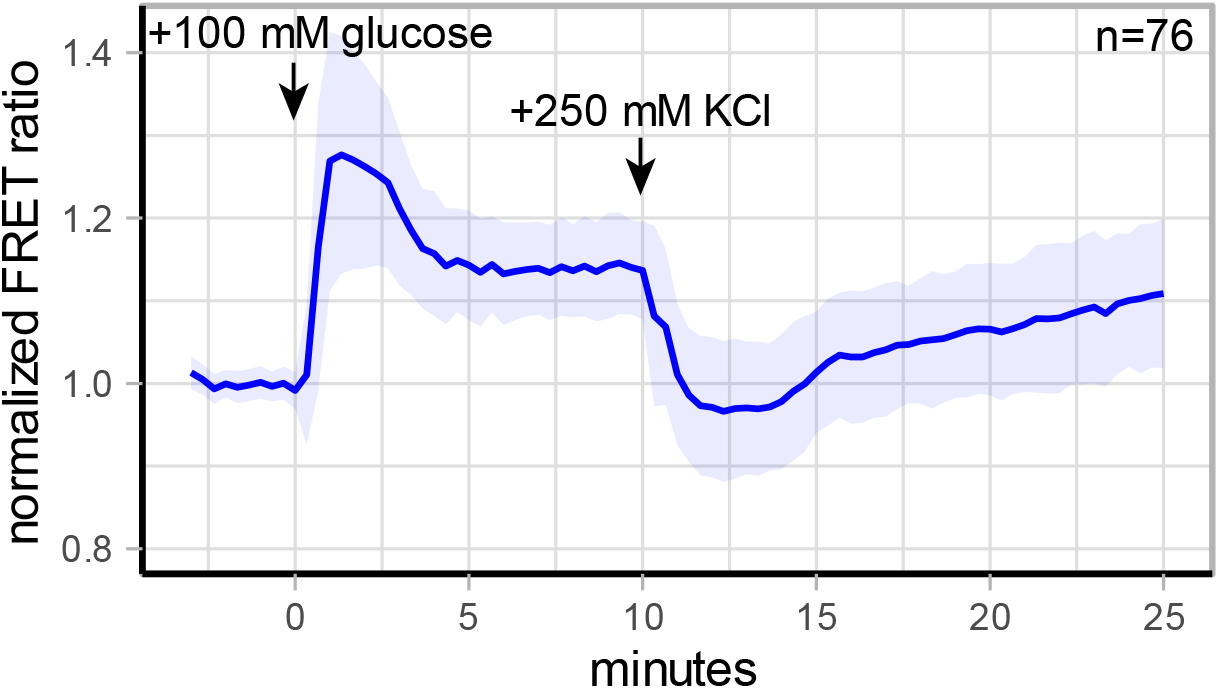
yEPAC FRET responses during salt stress. W303-1A WT cells grown on 1% ethanol were pulsed with 100 mM glucose at 0 minutes and with 250 mM KCl at 10 minutes (indicated with arrows). Lines show mean FRET ratios (corrected for the yEPAC-R279L response), normalized to the baseline, shaded areas indicate SD.

**Figure S6.**
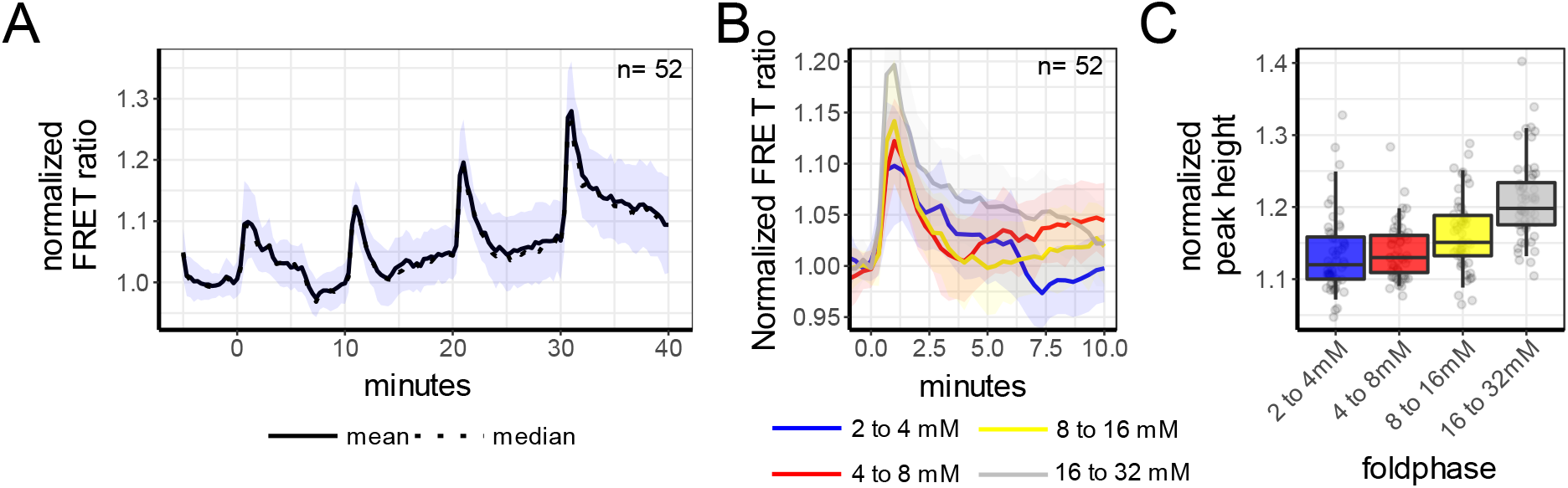
Weber-Fechner law of the same cells in time, with 10 minutes periods in between. A) W303-1A cells expressing yEPAC were pre-incubated on 2 mM glucose. Afterwards, cells were transitioned to 4 mM glucose at t=0 minutes and transitioned again to 8 mM at t=25 minutes. Lines show population mean response, normalized to the first 5 minutes, shaded areas indicate SD. B) Responses of the transitions performed in graph A, normalized to the last 3 frames before each transition. Colour indicates the transition. C) Boxplot of the peak heights of each cells for each transition. Dots show single-cell peak heights, boxes indicate median with quartiles, whiskers indicate the 0.05–0.95 fraction of the datapoints. D) Relation of normalized peak heights of single-cells between the first and second transitions. Dots represents single-cells.

**Figure S7.**
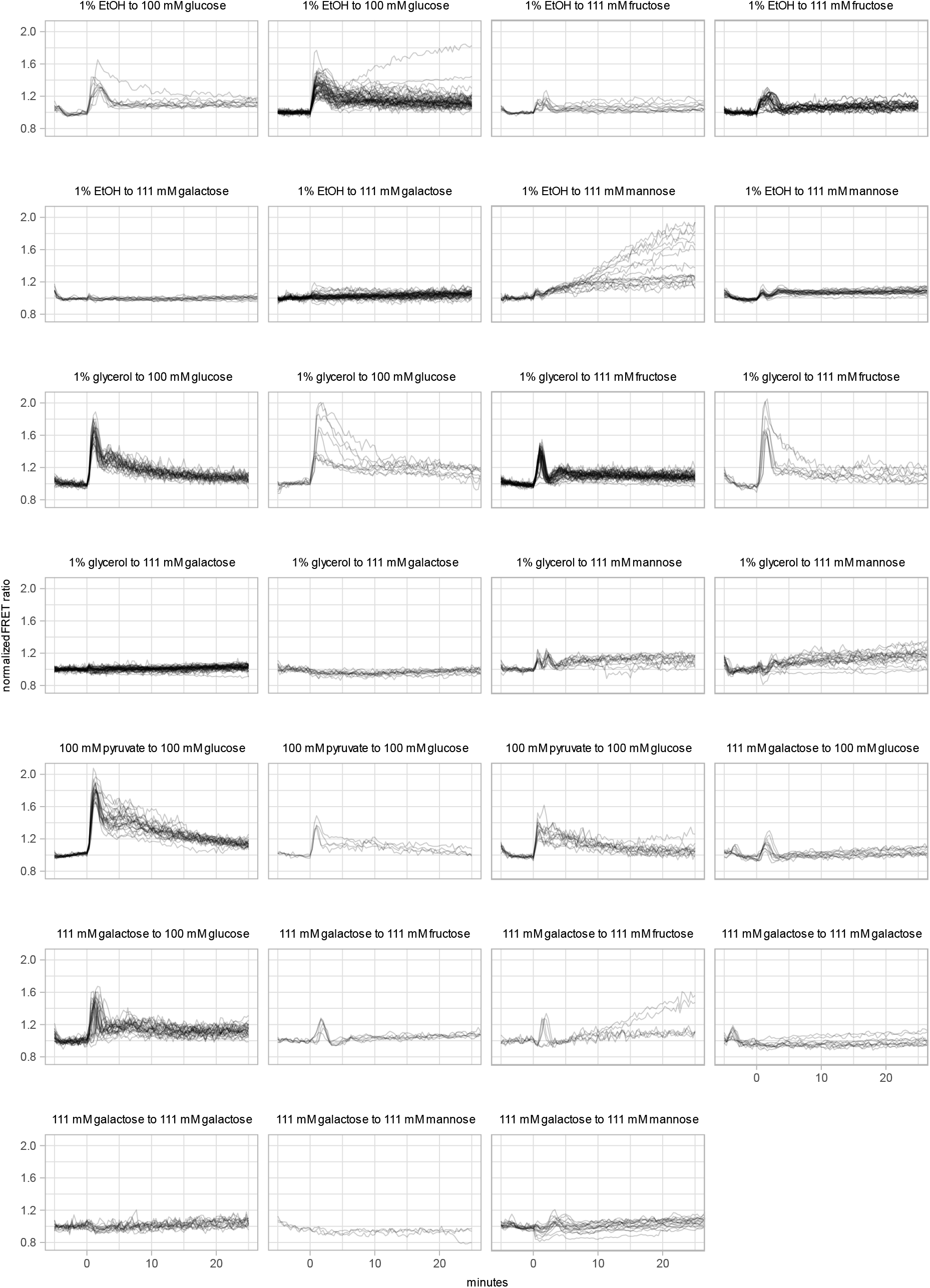
Single-cell yEPAC FRET responses of various carbon transitions. Cells growing on various non-fermentative carbon sources were pulsed with various sugars (transitions depicted above each panel) at 0 minutes. Lines show baseline-normalized FRET ratios of each cell.

**Figure S8.**
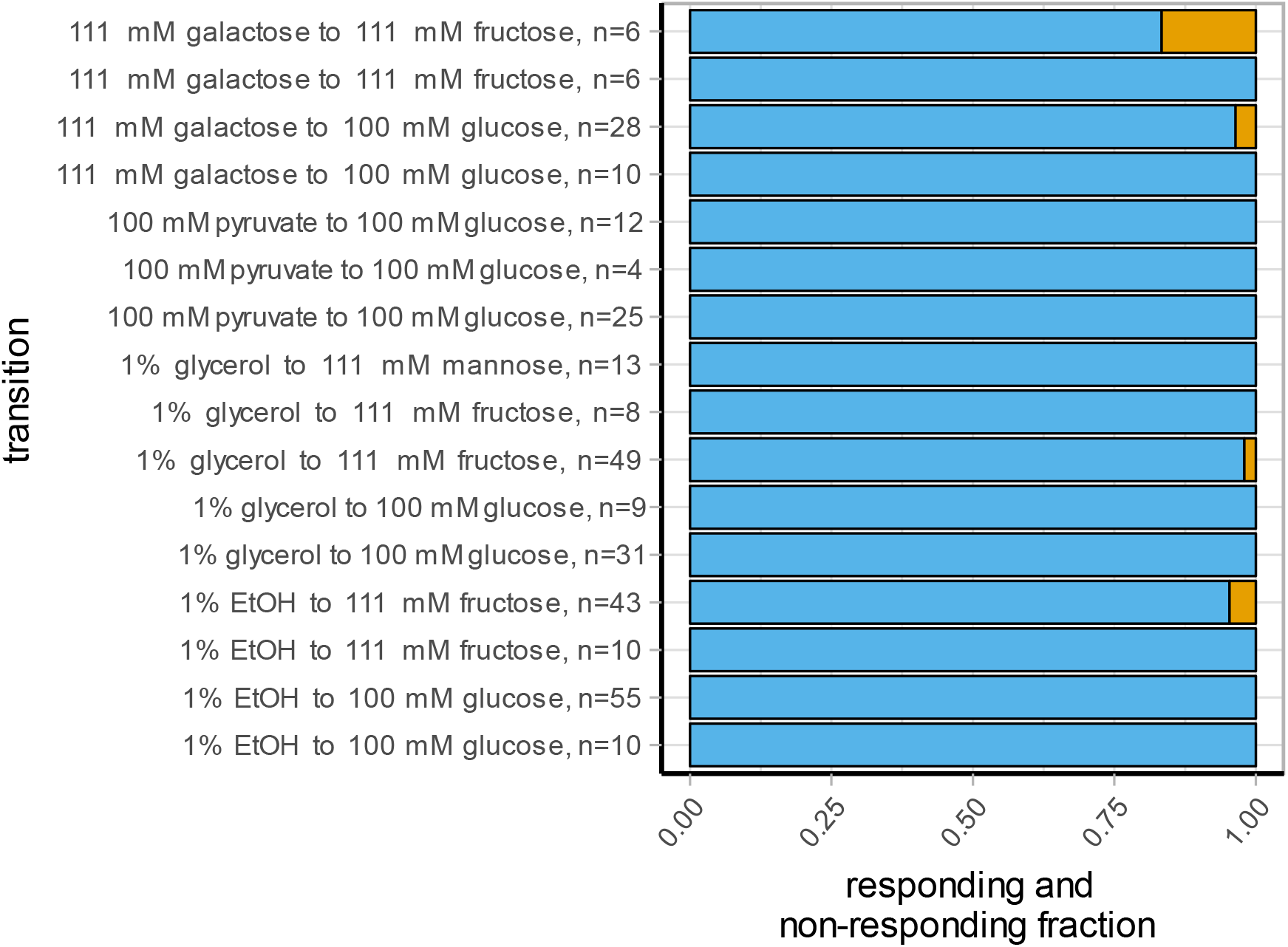
Fraction of responding (blue bars) and non-responding (orange bars) for the various glucose and fructose transitions. Only transitions that showed a population-average peak height of at least 1.15 were used. Cells were considered as responding when they showed a peak height of at least 30% of the mean peak value for the specific transition.

**Figure S9.**
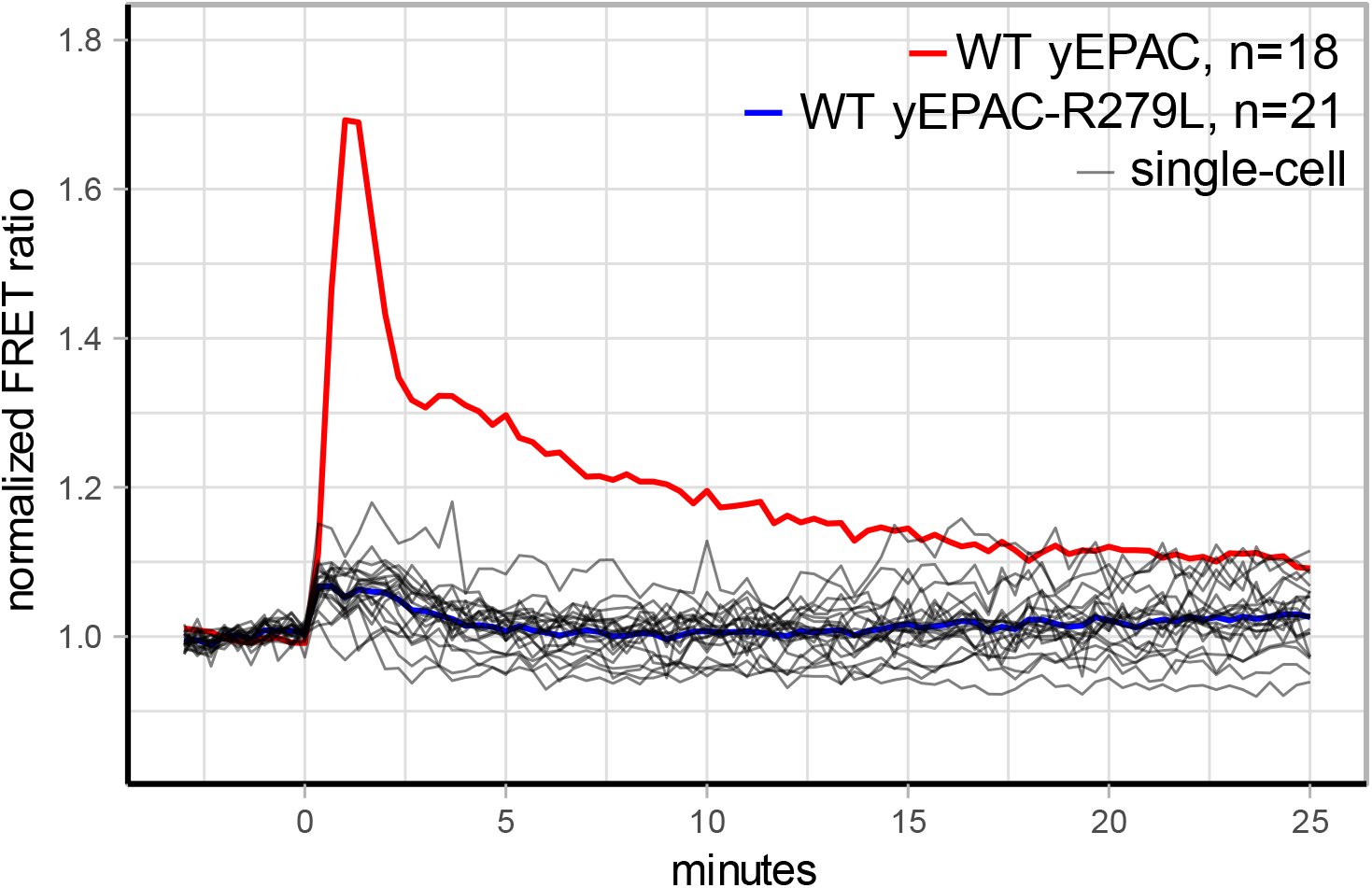
Long-term response of yEPAC-R279L after a transition from 1% EtOH to 100 mM glucose at t=0 minutes. The same data from figure 1 is depicted here with longer time scales. Thick coloured lines show mean response either yEPAC or yEPAC-R279L. Grey lines show single-cell responses, normalized to the baseline.

